# Molecular inflection points in the aging human brain

**DOI:** 10.1101/455105

**Authors:** Eden Deng

## Abstract

The onset of neurodegenerative diseases has been associated with age-dependent changes of gene expression in the brain. Research on age-dependent genes commonly assumes gradual linear relationships between gene expression and aging, failing to identify sudden changes in gene expression that may provide insight into the aging process and its role in neurodegenerative diseases. Here, a piecewise linear regression model is proposed to identify critical inflection points at which aging mechanisms may accelerate. Age-related gene expression data from tissue of four regions of the human brain (frontal cortex, hippocampus, putamen, substantia nigra) and human microglia were analyzed for inflection points. The best piecewise model represented stable expression during younger ages followed by an increase or decrease with age after the inflection point. In brain tissue, genes showing inflection points in expression pattern were enriched for gene ontology terms involved in neurodegenerative diseases. The hippocampus showed the highest proportion of genes with inflection points and the lowest variability in inflection points. These findings suggest that inflection points in gene expression may be used to characterize the aging process in the human brain and may help identify markers for the onset of neurodegenerative diseases.

## INTRODUCTION

The process of aging is a complex, dynamic phenomenon that influences the onset of many diseases. It is the greatest risk factor for neurodegenerative disorders such as Alzheimer’s and Parkinson’s, which affect nearly 7 million individuals in the United States alone [1, 2]. Although symptoms of the normal aging process do include slight cognitive decline, patients of neurodegenerative diseases experience accelerated decline and debilitating changes in behavior and brain function caused by neuronal death [3]. Individual brain regions are each uniquely affected by both aging and disease [4, 5]. Diagnosis of diseases has been associated with regional changes of gene expression in the aging brain, indicating the importance of research on changes in gene expression as the link between aging and disease.

Research on age-dependent genes commonly assumes a gradual linear relationship between gene expression and aging [6, 7]. However, this assumption can fail to capture the complexities of the aging process [8]. As the symptoms of age-related neurodegenerative diseases tend to occur at certain age, it is likely that underlying molecular changes can accelerate corresponding to age. Therefore, identifying sudden changes in age-dependent gene expression may provide insight into the aging process and its role in neurodegenerative diseases, which occur predominantly in the later stages of life.

Microglia play a unique and critical role in maintaining homeostasis in the central nervous system [9]. In an inactive state, they are constantly surveying and monitoring their local environment via dynamic processes. They are responsible for synaptic pruning, neuronal plasticity, clearance of cellular debris, and cell programmed death [10, 11]. However, in neurodegenerative diseases, microglia are activated to an inflammatory state and gather around deposits of misfolded protein, resulting in impairment of microglial function [12]. Because gene expression data from microglia is limited, the age-associated changes of microglia and their contributions to the onset of neurodegenerative diseases are poorly understood. Although recent studies of the microglia transcriptome and its gene signature further support the role of microglia in disease [13, 14], the cell type’s age-dependent gene expression profile remains unclear.

Because of microglia’s role in disease, it is possible that the gene expression patterns of disease-related brain regions reflect the gene expression patterns of microglia. Investigating age-dependent microglial gene expression in the context of aging in the brain cortex may provide clues regarding the nature of late-onset neurodegenerative diseases. Because of the role of microglia in brain aging, we are particularly interested in the age-dependent gene expression changes in microglia.

## METHODS

### 1. Data Sources and Processing

Brain tissue gene expression data was obtained from the United Kingdom Brain Expression Consortium (UKBEC), accessible through Gene Expression Omnibus GSE60862. Data quality and reliability was confirmed in Trabzuni *et al.* [15]. Affymetrix probes were mapped to official gene symbols using the Bioconductor package “huex10sttranscriptcluster.db”. Four subsets of samples from different brain regions were generated from the UKBEC data set: frontal cortex (n=127), hippocampus (n=122), putamen (n=129), and substantia nigra (n=101). Multiple probe sets from the same gene were summarized by mean, and genes with little variability (sd<0.2 in log2 scale) within each subset were filtered out.

Microglia gene expression data was obtained through Galatro *et al.* [14] in which RNA sequencing was performed on microglia isolated from human postmortem parietal cortex brain tissue. The data set, accessible through Gene Expression Omnibus GSE99074, contained 39 microglia samples from donors between 34 and 102 years of age and 16 corresponding samples of parietal cortex tissue to serve as a control group. As described in Galatro *et al.*, the data were normalized with voom and annotated using BioMart. Analysis for batch effects due to age, gender, or country of origin was performed using the *limma* package and PCA. Genes with low variation (sd<0.5) were removed. An outlier sample (SPM31) was removed.

### 2. Differential Expression Analysis

For all data sets, to identify genes associated with age, a linear regression model was fit between gene expression and age using the function *lmFit* from *limma*. The differential expression criteria for age-dependent genes was set to a p value<0.05. Enrichment values between the age-dependent genes of two different data sets were calculated by dividing the multiple of both fractions of age-dependent genes (expected by chance) in the human genome from the observed number of overlapping age-dependent genes. A score of 2, for instance, means that the observed overlap is twice the expected overlap due to chance.

### 3. Inflection Point Identification

Piecewise linear models, describing segmented linear models with inflection points, is one of the simplest nonlinear functions. We are interested in the function where an inflection point occurs in an age-dependent manner. We hypothesize that genes associated with the late-onset of neurodegenerative diseases will have a stable expression during younger ages and start to increase or decrease with age after a certain age. To identify these genes with inflection points, we started with the top 1000 age-associated genes identified in section 2. We fit a linear model of gene expression with age

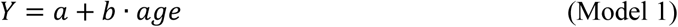

where we did not explicitly include the error term for simplicity. Then we fit three inflection models with an unknown inflection point *k* in age (ModelA, ModelB, and ModelC)

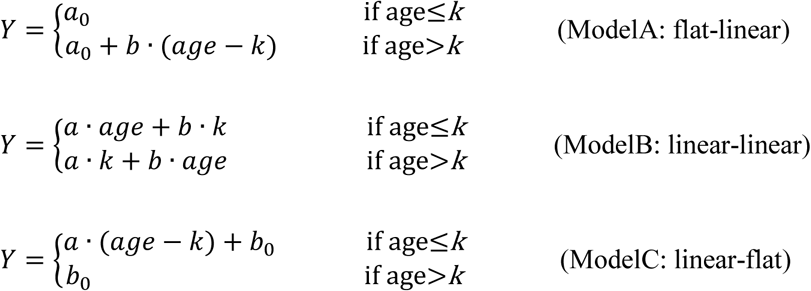

ModelA presents an early stable or flat gene expression followed by an onset of age-dependent change at some age *k*. ModelB presents a change in slope of the age-dependent relationship. ModelC presents an initially age-dependent gene expression followed by stable or flat expression. The three models represent three possible patterns of a piecewise linear relationship between age and gene expression. R package *segmented* was used to estimate the unknown inflection point *k* and test the significance of the inflection point model by comparing Model1 with ModelA, ModelB, or ModelC. An F-test was used to calculate the p values when comparing the two models and their goodness of fit. The criteria for a significant age inflection point model was set as p value<0.05. Multiple comparison adjusted p values were not used due to the small sample sizes. Because the UKBEC data contained a larger sample size, outlier samples with values more than 3 standard deviations from the mean were removed. Enrichment for gene ontology (GO) terms of genes exhibiting an age inflection point was performed with DAVID [16].

All analyses were conducted in R (version 3.3.3).

## RESULTS

### Evaluation of Inflection Models

The fits of inflection models (ModelA, ModelB, ModelC) were compared to those of the standard linear regression model (Model1). Of the three inflection models tested, 89% of all genes with inflection points fit (p value<0.05) ModelA, the flat-linear model, indicating a stable gene expression followed by an onset of age-dependent expression is the most likely pattern for the nonlinear relationship we evaluated. 28.2% of all age-dependent genes were better fit to ModelA than the linear model. The analyses showed that ModelB and ModelC were less representative of brain aging patterns.

### Inflection Points in UKBEC Brain Tissue

Shown in Figure 1, the inflection points of each region (identified through ModelA) showed peaks in their distribution. The mean inflection point in the frontal cortex, hippocampus, putamen, and substantia nigra was 65.4, 52.9, 64.5, and 61.5 years of age, respectively. The standard deviation for the four brain regions were 8.1, 5.5, 7.4, and 6.6, respectively. On average, 78% of the genes showing a inflection point were downregulated during aging. The genes with inflection points in each brain region were enriched for cellular dysfunctions related to the late-onset of neurodegenerative diseases (Figure 2). In the frontal cortex, genes involved in the MAPK pathway (CCL2, VEGFA) were downregulated. Genes involved in the restriction of neuro-pathogenic protein aggregation (CCT3, CASP2) were dysregulated in the aging hippocampus. In both the putamen and the substantia nigra, genes associated with the prevention of oxidative stress showed decreased expression.

**Fig1.**
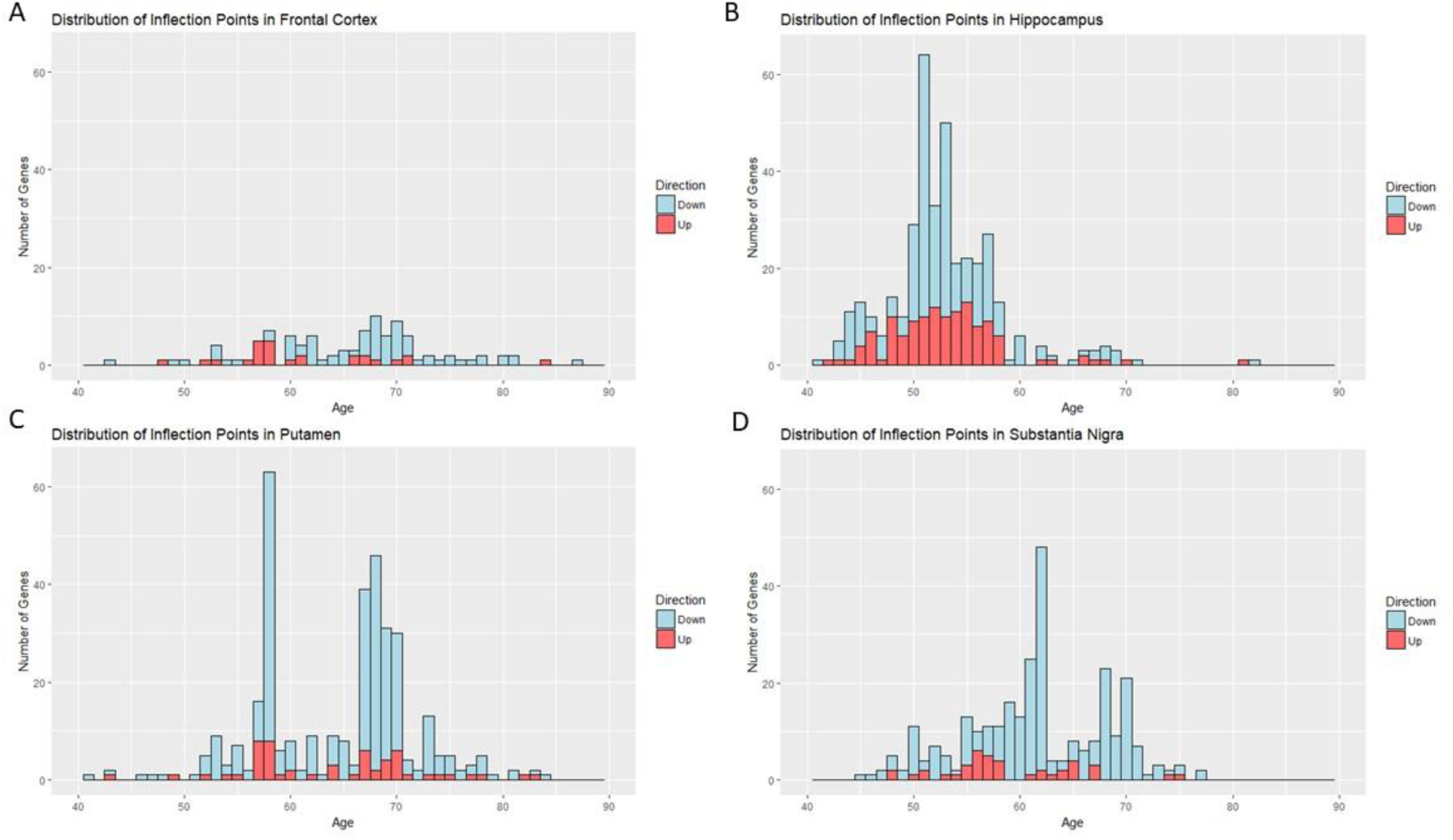
Distribution of inflection points (ModelA) in UKBEC data set. The blue and red bars show the number of age-dependent genes with down-regulation and up-regulation, respectively, after inflection points. A) The frontal cortex showed the lowest proportion of inflection points (10.3%, n=103). B) The hippocampus showed the highest proportion of inflection points (38.2%, n=383). C) Putamen (35.3%, n=353). D) Substantia nigra (28.8%, n=288).

**Fig2.**
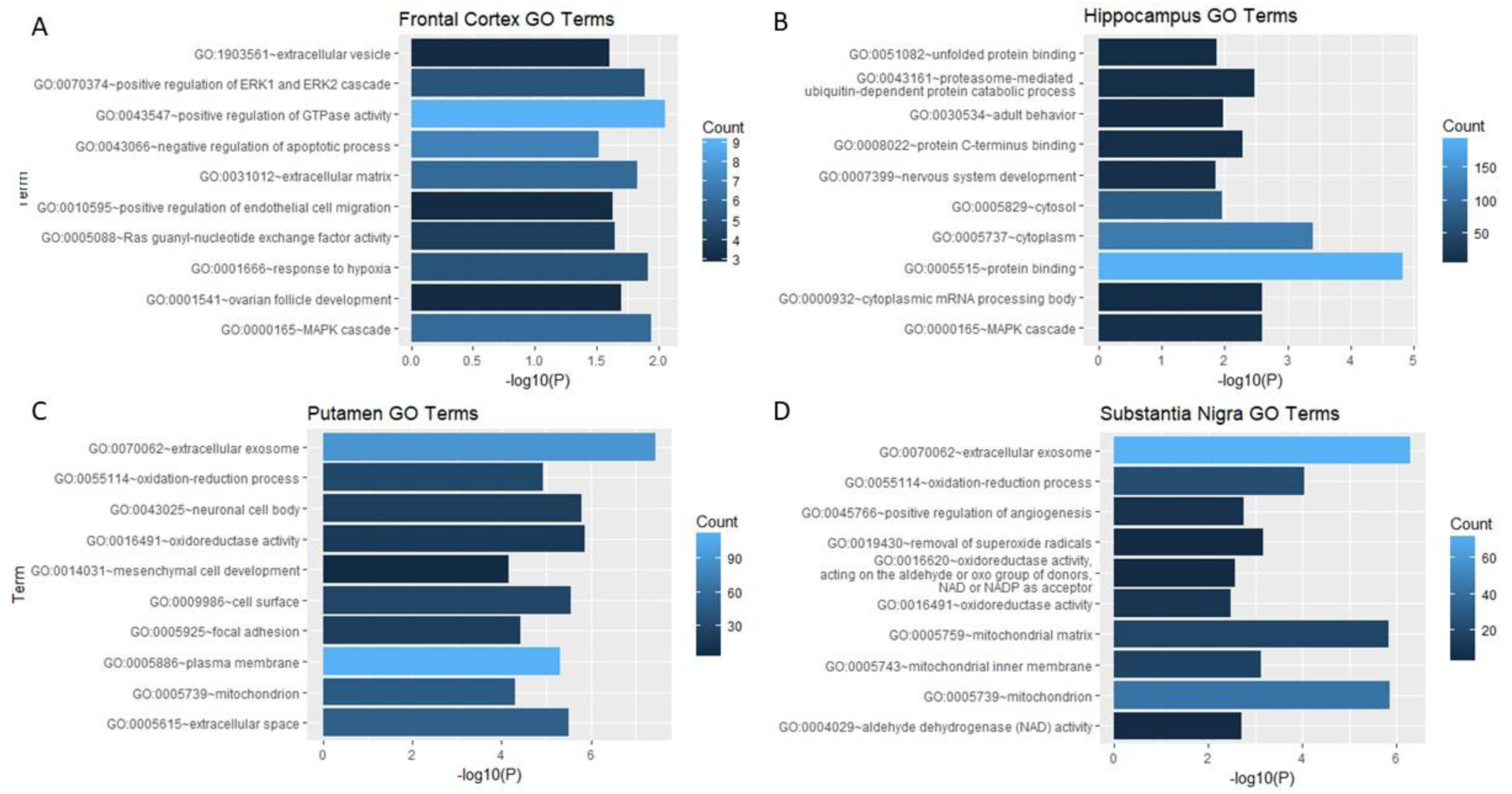
Functions of genes with inflection points (UKBEC). Top 10 enriched GO terms are displayed, where the lengths of the bars represent statistical significance (-logP) and the color corresponds to gene count. A) Frontal cortex. B) Hippocampus. C) Putamen. D) Substantia nigra.

### Inflection Points in Parietal Microglia

Heatmaps revealed differentially expressed genes based on a linear regression model between gene expression and age (Figure 3). 84 microglia genes (p value<0.001) showed a separation in gene expression between older and younger individuals, as seen in Galatro *et al.* 52 genes were upregulated during aging, and 32 were downregulated. Shown on the right are the 73 differentially expressed genes in cortex samples (p value<0.001), two of which overlap with the age-associated microglial genes. As shown in Figure 4, 58 microglial genes (p value<0.05) exhibit inflection points at which gene expression transitions from a constant level of expression to an age-dependent pattern of expression (ModelA). The distribution of inflection points peaks near age 65 (mean=67.2, sd=5.6). 74% of the genes shown were downregulated during aging. Notable genes with reduced expression in microglia during aging and an inflection point in gene expression include regulators of calcium ion transport and lipid metabolism. For example, CACNB2 and PLCG2 showed inflection points at 66.9 and 64.2 years of age, respectively, after which gene expression begins to decrease with age.

**Fig3.**
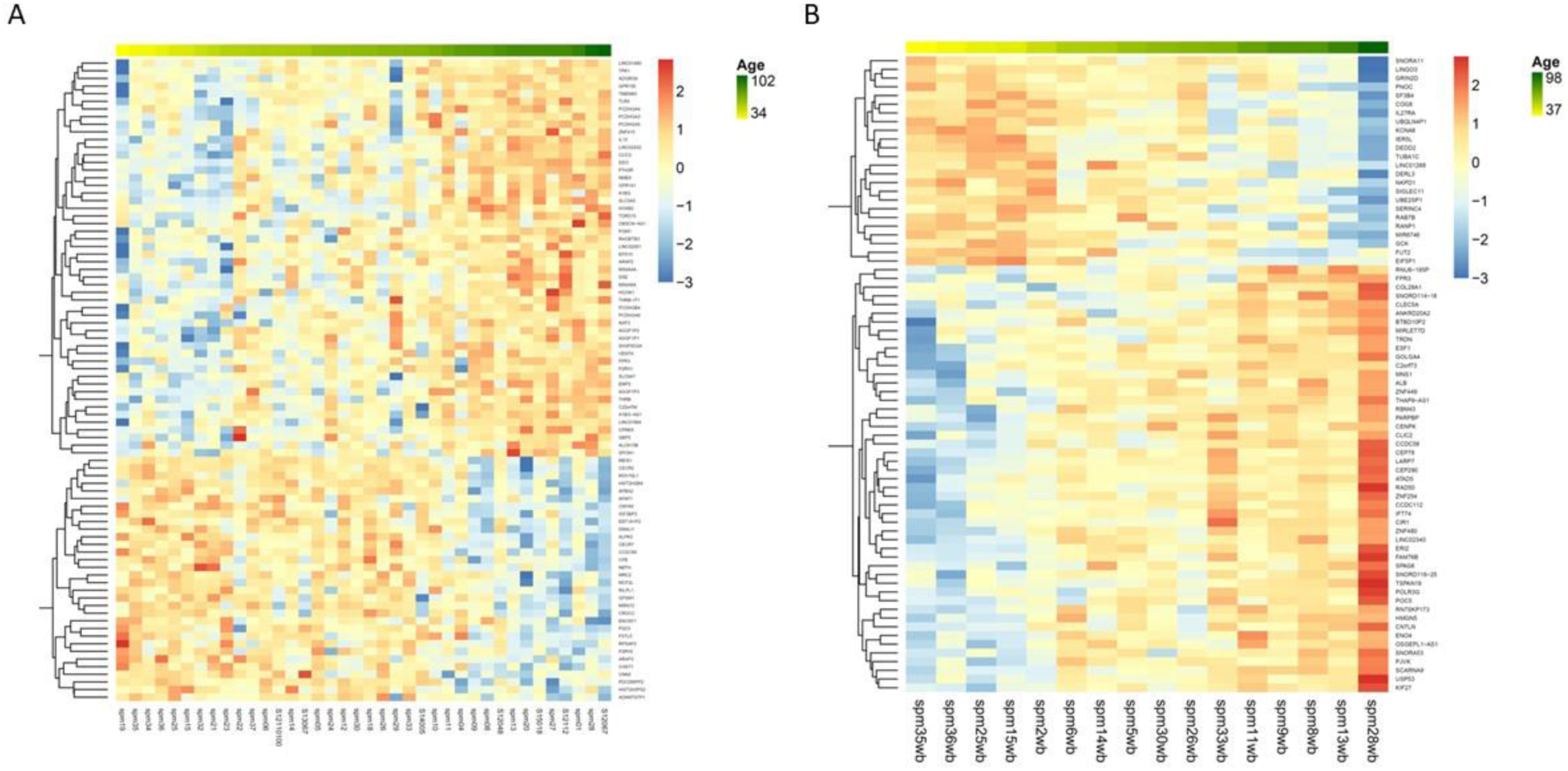
Heatmaps of age-dependent gene expression from Galatro data set. Standardized expression heatmaps of differentially expressed genes (p<0.001) showed age-dependent clustering in samples. A) Microglia from parietal cortex. B) Parietal cortex tissue.

**Fig4.**
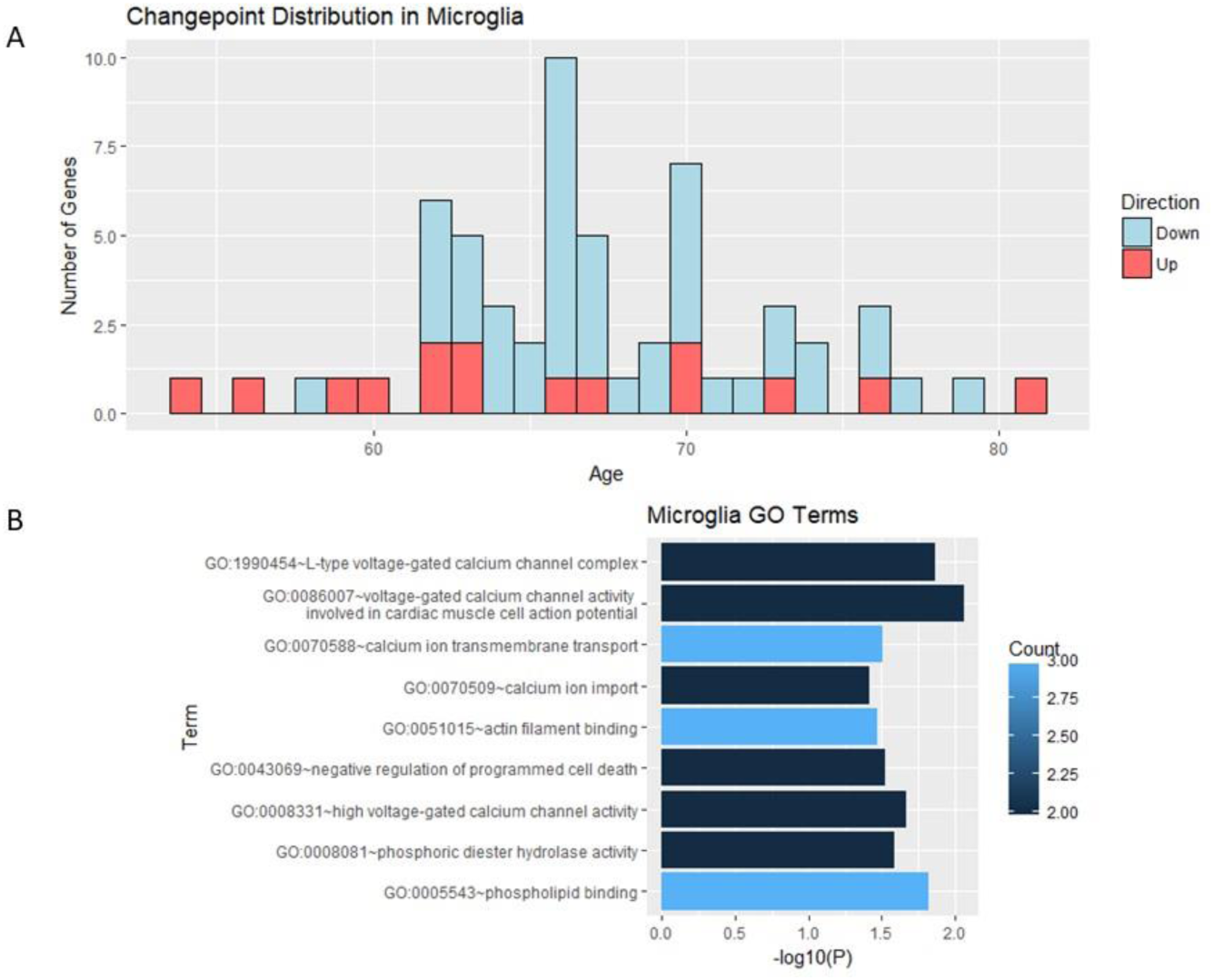
Distribution and functions of genes with inflection points in microglia. A) The blue and red bars show the number of age-dependent genes with down-regulation and up-regulation, respectively, after inflection points. Microglia showed 5.9% of age-dependent genes with inflection points (n=59). B) Top 10 enriched GO terms in microglial genes with inflection points, where the lengths of the bars represent statistical significance (-logP) and the color corresponds to gene count.

### Intersection Between Data Sets

The age-dependent in microglia were not enriched in other brain regions, and the age-dependent genes of different brain regions also showed little overlap. The substantia nigra and putamen regions showed the most overlap (enrichment=4.62). The hippocampus showed a more similar age-dependent profile with the putamen and substantia nigra (enrichment=2.52 and enrichment=4.16, respectively). For all data sets, there was no significant enrichment between the inflection point genes.

**Table 1.** Overlap of age-dependent genes. The age-correlated genes (p value<0.05) between brain regions and microglia showed very little overlap (mean enrichment=1.00). The genes from each brain region showed more moderate overlap with each other, particularly the substantia nigra and putamen, where the enrichment value was 4.62.

|  | Frontal cortex | Hippocampus | Putamen | Substantia nigra |
| --- | --- | --- | --- | --- |
| Microglia | 0.86 | 0.9 | 1.14 | 1.1 |
| Frontal cortex |  | 1.62 | 2.26 | 2 |
| Hippocampus |  |  | 2.52 | 4.16 |
| Putamen |  |  |  | 4.62 |

## DISCUSSION

### Inflection Models of Aging

In the evaluation of patterns of nonlinear gene expression in the aging human brain, the three inflection models tested revealed a significant number of genes exhibiting inflection points during aging in all data sets. The best model (ModelA) represented 28.2% of all age-dependent genes. Although linear models remain most representative of brain aging, ModelA characterizes a notable aging pattern in which the initial stability of molecular processes are disrupted through the onset of age-dependent change in gene expression.

### Frontal Cortex

Among all brain regions, aging in the frontal cortex is the most thoroughly researched and has been strongly linked to cognitive decline. The frontal cortex is generally considered to undergo age-related changes before other cortical regions, as prior studies [17, 18] support the frontal lobe hypothesis of aging. Here, very few genes with inflection points were identified in the frontal cortex relative to other brain regions. This raises the question of whether most molecular changes in the frontal lobes occur before the scope of this study at an earlier age. The genes with inflection points that were identified in the frontal cortex were enriched for MAPK signaling and GTPase regulation. The MAPK pathway regulates a variety of neuronal functions (proliferation, differentiation, survival, and death) [19, 20], and Rho GTPases are regulators of the cytoskeleton and mediate diverse neuronal processes [21]. Changes in the regulation of both MAPK signaling and GTPase activity have been implicated in the development of many neurodegenerative diseases [22, 23].

### Hippocampus

Dementia is the most prominent symptom of Alzheimer’s, resulting from shrinkage of the hippocampus [24]. The high proportion and lower variability of genes with inflection points (38.2% of age-dependent genes fit ModelA, sd=5.5) in the hippocampus points towards a significant age-dependent change. Many of the down-regulated genes with inflection points are involved in the prevention of neuro-pathogenic protein aggregation. Notably, the downregulation of CCT3 and upregulation of CASP2 are associated with the aggregation of the toxic proteins found in Alzheimer’s disease, Parkinson’s disease, and Huntington’s disease [25, 26]. Expression changes in these genes were centered around age 53,preceding the most common diagnosis age of late-onset Alzheimer’s [27]. Thus, our results are supported by well-established findings. These hippocampal expression changes may potentially serve as precursors to late-onset Alzheimer’s.

### Putamen & Substantia Nigra

The putamen and substantia nigra are both components of the basal ganglia and are involved in the neurotransmission of dopamine. The substantia nigra region is associated with Parkinson’s disease [28], a disorder resulting from the degeneration of dopaminergic neurons. Its pathology has also been linked to the progression of Alzheimer’s disease [29]. A notable number of genes with inflection points were identified in both the putamen and substantia nigra (35% and 29% of differentially expressed genes, respectively), suggesting a significant age-dependent change. Both regions exhibited inflection points in genes associated with the prevention of oxidative stress, which, in Parkinson’s disease, results in mitochondrial dysfunction and an increase in reactive oxygen species, leading to cell death [30]. Both regions also had a similar bimodal shape in their inflection point distributions, although the causes of such a shape are unclear. The similarities between the two distributions, however, support the close association between the putamen and substantia nigra as well as their role in neurodegenerative diseases.

### Microglia

Microglia showed very few genes with inflection points, suggesting its pattern of gene expression is either stable with age or consistently linear with age. The microglial genes with inflection points were enriched for functions of calcium ion transport and lipid metabolism. As subunits of voltage-gated calcium channels, CACNA1D and CACNB2 contribute to cellular regulation of calcium ion signaling [31, 32]. Microglial function has been found to be dependent on intracellular calcium ion signaling [33, 34], and in neurodegenerative diseases, altered microglial function and dysregulation of cytokine release contributes to degeneration of neurons [11]. Previous research [35] also suggests that many neurodegenerative disorders are associated with microglia-mediated dysregulation of lipid metabolism.

### Comparisons Across Data Sets

Analysis of both brain tissue and microglia gene expression data showed that the number of inflection points peaked in the 50s or 60s (total mean=60.2, total sd=8.5). On average, 75% of these genes with inflection points were downregulated during aging. The consistent peaks in inflection point distribution and the strong down-regulation of inflection point genes raise the question of whether a group of transcriptional repressors in each brain region become active once a critical age threshold is reached. Microglial age-dependent genes and age-dependent genes from the frontal cortex, hippocampus, putamen, and substantia nigra shared little overlap (enrichment≤1.14), although this may have been due to microglia samples being from the parietal cortex. Prior studies [4, 5] have observed that different brain regions show different patterns of response to aging, so shared aging processes across the brain may not be detected. Here, the frontal cortex, hippocampus, basal ganglia regions each showed different responsiveness to age-related changes and different molecular changes.

### Future Steps

Molecular inflection points in the aging human brain, particularly the hippocampus, may serve as markers for the onset of neurodegenerative diseases and targets for preventative treatment. However, the role of age-related inflection points in disease must be further studied and confirmed as predictors of disease. Mapping the transcriptional regulation networks of these inflection points may provide a better understanding of the relationships between genes with inflection points.

## SUPPLEMENTARY INFORMATION

S1 Table. Differentially expressed genes (UKBEC)

Gene expression in the frontal cortex, hippocampus, putamen, and substantia nigra versus aging.

S2 Table. Inflection points from ModelA (UKBEC and Galatro)

Genes with inflection points in the frontal cortex, hippocampus, putamen, and substantia nigra cortical regions and microglia.

S3 Table. GO terms for genes with inflection points

Gene ontology terms associated with the genes with inflection points in the frontal cortex, hippocampus, putamen, and substantia nigra cortical regions and microglia.

